# Expression and Purification of Full-Length hnRNPA2B1 for Biophysical Characterization of Liquid-Liquid Phase Separation

**DOI:** 10.1101/2025.07.10.664112

**Authors:** Luis Fernando Durán-Armenta, Attila Meszaros, Julia Malo Pueyo, Steven Janvier, Tamas Lazar, Dominique Maes, Remy Loris, Peter Tompa

## Abstract

Heterogeneous nuclear ribonucleoprotein A2/B1 (hnRNPA2B1) is a multifunctional RNA-binding protein involved in RNA maturation and mRNA transport. It has recently been shown to undergo liquid-liquid phase separation (LLPS), contributing to the assembly of membraneless organelles. Moreover, dysregulation of LLPS is associated with the formation of pathogenic protein aggregates, in which hnRNPA2B1 is frequently found. Despite its biological and pathological relevance, studies on the full-length protein remain limited due to its intrinsically disordered, low-complexity domain, which renders hnRNPA2B1 highly aggregation-prone and difficult to purify. In this study, we report the successful expression and purification of full-length hnRNPA2B1 with high purity and minimal nucleic acid contamination. By optimizing buffer conditions, specifically ionic strength and pH, we maintain the protein in solution following cleavage of its solubility tag. Preliminary *in vitro* characterization under near-physiological conditions reveals that purified hnRNPA2B1 undergoes LLPS, forming dynamic liquid-like droplets that grow and mature into amorphous aggregates. Our approach provides a robust method for purifying hnRNPA2B1 suitable for LLPS and aggregation studies. This strategy may also be useful to purify other aggregation-prone, intrinsically disordered proteins.

## 1. Introduction

Heterogeneous nuclear ribonucleoprotein A2/B1 (hnRNPA2B1) is an RNA-binding protein that plays key roles in multiple aspects of RNA metabolism, including pre-mRNA splicing [1,2], mRNA transport [3], and mRNA stability [4]. In recent years, several members of the hnRNP family, including hnRNPA2B1, have gained attention in the context of biomolecular condensates. hnRNPA2B1 undergoes liquid-liquid phase separation (LLPS) in eukaryotic cells, contributing to the formation of membraneless organelles (MLOs) such as stress granules [5] and other ribonucleoprotein granules [6,7]. LLPS is primarily driven by its C-terminal intrinsically disordered low-complexity domain (LCD), which is enriched in glycine, asparagine, tyrosine, and arginine residues. These residues enable multivalent, transient homotypic and heterotypic interactions with other phase-separating proteins [6] and nucleic acids [6,8]. This region also contains low-complexity, amyloid-like, reversible kinked segments (LARKS), which facilitate self-association [5] and the formation of reversible hydrogels and amyloid-like assemblies [9]. Growing evidence suggests that LCD-mediated phase separation represents a mechanistic link between MLO assembly and fibrillar protein pathology [10,11]. Aging, persistent stress, and disease-linked mutations can alter condensate dynamics, promoting their maturation into pathogenic, irreversible aggregates [5,6]. Notably, several hnRNPs, including hnRNPA2B1, have been identified in the proteome of both stress granules and pathological inclusions in neurodegenerative diseases such as amyotrophic lateral sclerosis (ALS) and Multisystem Proteinopathy (MSP) [5,11–13].

Given the importance of hnRNPA2B1 in MLO assembly and disease, *in vitro* studies have examined the phase behavior [6,14], the effects of disease-linked mutations on aggregation [5,6], and the structural properties of amyloid fibers [9]. However, these studies mainly employ isolated domains, either the globular RNA Recognition Motifs (RRMs) [15] or LCD chimeras fused to solubility tags [5,6] or fluorescent proteins [9]. Studies on full-length hnRNPA2B1 are scarce, largely due to the lack of established protocols for its expression and purification. hnRNPs and other intrinsically disordered proteins are challenging to purify due to their limited solubility, high aggregation propensity, and susceptibility to degradation. Nevertheless, studying full-length hnRNPA2B1 is essential to understanding its native behavior, including potential interactions between its RRMs and LCD. Such interplay has been shown to modulate phase separation in the closely related paralog hnRNPA1 [16].

Other challenging full-length hnRNPs, such as hnRNPA1 [17], hnRNPDL [18], and TDP-43 [19], have been successfully purified using solubility tags and denaturants. Whereas these strategies may increase protein solubility and yield, they may also introduce artifacts in LLPS and aggregation assays, which are highly sensitive to experimental conditions such as temperature, nucleic acid contamination, pH [14], and ionic strength [14,18]. Residual denaturants, proteases, uncleaved protein, and other impurities may alter condensate behavior and lead to different kinetic paths [14].

In this study, we report a novel purification protocol to obtain pure, soluble full-length hnRNPA2B1 without denaturants or residual solubility tags. Although we employ a SUMO-hnRNPA2B1 fusion for recombinant expression and early purification steps, we successfully maintain the protein in solution following tag removal by carefully optimizing the ionic strength and pH [14]. The identity and purity of the purified protein were confirmed by mass spectrometry, analytical size exclusion chromatography, and mass photometry. Furthermore, we demonstrate that purified hnRNPA2B1 undergoes LLPS *in vitro*, forming spherical, liquid-like droplets that mature into amorphous aggregates.

This approach provides a robust strategy for purifying hnRNPA2B1 and is intended to serve as a framework for purifying other challenging proteins from this family, including full-length hnRNPs and other intrinsically disordered proteins, under native-like conditions. Our approach will facilitate further functional and structural studies of hnRNPA2B1, particularly in the context of phase separation and protein aggregation.

## 2. Materials and Methods

### Bioinformatics

Disorder prediction was performed using AIUPred [20] and structural dynamics/rigidity was assessed using DynaMine [21,22], both with default parameters via their respective web servers. The CIDER web tool [23] was used to predict the theoretical collapse of the polypeptide chain. Protein aggregation propensity was predicted by AGMATA [24] in single sequence mode, while homotypic LLPS propensity was predicted by FuzDrop [25]. The predicted AlphaFold2 model [26] was downloaded from the AlphaFold Protein Structure Database [27]. All analyses were conducted using the canonical sequence of hnRNPA2B1 (UniProt accession: P22626) [28].

### Vector and Protein Expression

The prokaryotic expression vector pET-22b (+) encoding SUMO-hnRNPA2B1 was purchased from GenScript. This plasmid expresses human hnRPNA2B1 with a C-terminal hexahistidine SUMO tag. A TEV protease cleavage site (ENLYFQS) was included between SUMO and hnRNPA2B1 to enable tag removal.

The construct was transformed into chemically competent *Escherichia coli* BL21 STAR™ (DE3) cells via heat shock. Transformed colonies were selected on LB-agar plates supplemented with 100 µg/mL carbenicillin. A single colony was used to inoculate a 50 mL LB preculture with carbenicillin and incubated overnight at 37ºC. This preculture was diluted 1:100 into 1 L of Terrific Broth (TB) with carbenicillin and grown at 37ºC until OD_600_ reached 1.0-1.2. Protein expression was induced with 1 mM isopropyl β-D-1-thiogalactopyranoside (IPTG), and cultures were incubated for an additional 4 hours at 37 ºC before harvesting by centrifugation at 5000 x g for 15 minutes.

### Electrophoresis and immunoblotting

Protein samples were mixed with SDS Loading Dye and separated on 4–15% Mini-PROTEAN® TGX™ Precast Protein Gels (Bio-Rad) using the PageRuler™ Prestained Protein Ladder (Thermo Scientific) as a molecular weight marker. Gels were stained using PageBlue™ Protein Staining Solution (Thermo Scientific).

Proteins were transferred to a Trans-Blot® Turbo™ Mini 0.2 µm PVDF membrane (Bio-Rad) using the Trans-Blot® Turbo™ Transfer System. The membranes were blocked with 5% non-fat milk in TBS-T (Tris-buffered saline with 0.1% Tween-20) for 1 hour at room temperature, then incubated overnight at 4ºC with a mouse monoclonal anti-polyHistidine-Peroxidase antibody (Sigma-Aldrich, 1:1000 dilution). After washing with TBS-T, the membranes were developed with Pierce® ECL Western Blotting Substrate (Thermo Scientific) and imaged with an Amersham ImageQuant 800 GxP biomolecular imager (Cytiva).

### Protein Purification

A pellet was resuspended in 100 mL lysis buffer (50 mM HEPES, 1M NaCl, 10 mM imidazole, 0.5 mM TCEP, pH 8.0) supplemented with 0.1% Triton X, RNAse A, DNAse I, 0.1 mM PMSF, 0.5 mM benzamidine hydrochloride, and 2 cOmplete™ EDTA-free protease inhibitor tablets (Roche). Cells were lysed by sonication on ice using a VCX-70 Vibra-Cell™ (Sonics & Materials) for 10 minutes (5 s pulse on, 5 s pulse off, 70% amplitude). An additional 50 mL of lysis buffer was added before centrifugation (18,000 rpm, 4ºC, 1 h). The supernatant was filtered and loaded onto a 5 mL HisTrap™ HP column (GE Healthcare) equilibrated with wash buffer (50 mM HEPES, 1M NaCl, 10 mM imidazole, 0.5 mM TCEP, pH 8.0). Bound SUMO-hnRNPA2B1 was eluted with a linear gradient of wash buffer supplemented with 500 mM imidazole. Pooled fractions were desalted into Cleavage Buffer (50 mM Tris, 1M NaCl, 0.5 mM TCEP, pH 7.5), and 500 µL of TEV protease (5 mg/mL) were added for overnight incubation at room temperature. The solution was then loaded onto a second HisTrap™ HP column equilibrated with wash buffer without imidazole. The flow-through containing tag-free hnRNPA2B1 was collected. Selected fractions were then dialyzed into 10 mM CAPS, pH 11.0, aliquoted at 15 µM, and flash-frozen in liquid nitrogen for storage at -80ºC.

Protein concentration was determined using a Nanodrop™ One (Thermo Scientific). Theoretical extinction coefficients (*ε*) were calculated using ProtParam: *ε =* 41260 M^-1^ cm^-1^ for SUMO-hnRNPA2B1 and *ε =* 39770 M^-1^ cm^-1^ for hnRNPA2B1. Protein purity was assessed by SDS-PAGE and Nanodrop™ One. Pure samples showed an A260/A280 ratio of approximately 0.6, indicating minimal nucleic acid contamination.

### Mass Spectrometry

Matrix Assisted Laser Desorption Ionization – Time of Flight (MALDI-TOF) MS data was acquired on a Ultraflextreme enhanced MALDI TOF/TOF-MS system (Bruker Daltonics, Bremen, Germany) using FlexControl 3.4 acquisition software (Bruker).

For intact mass determinations, 1 µL of a 1:1:1 mixture of protein sample, 2.5 DHAP matrix (Bruker) and 2% *v/v* TFA was spotted in duplicate on an MTP ground steel plate. After crystallization, spectra were measured in the linear positive ion-mode within a mass range of 10 000 to 100 000 m/z. Up to 8000 shots were acquired with a laser repetition rate of 2000 Hz and 200 shots per raster spot. All acquisition methods were provided by the manufacturer and optimized and calibrated with in-house made calibration standards (10 to 44 kDa, 4 calibrants). The obtained spectra were analyzed and processed (peak picking, smoothing and baseline substraction) with FlexAnalysis 3.4 (Bruker). Finally, the average of the two sample spots was taken to obtain the measured mass.

For top-down sequencing (TDS) experiments to confirm the N- and C-terminus, 10 µL sample was mixed with 8M GuHCl, desalted with a C18 ZIPTIP (Millipore) and eluted with a 50 mg/mL SDHB solution (dissolved in 50:50 Acetonitrile:0.1% Trifluoroacetic acid). The eluent was diluted twofold with deionized water and 1.5 µL spotted on an MTP ground steel plate. All TDS experiments were measured in reflector and ion-positive mode within a range of 500 to 9000 Da with a by the manufacturer provided optimized acquisition method. Up to 12000 shots were manually acquired and the method used was optimized and calibrated with an in-house made protein (1-6 kDa). The obtained spectra were processed and analyzed with FlexAnalysis 3.4 (Bruker) and BioTools 3.2 (Bruker).

To further confirm the amino acid sequence a tryptic digest was subjected to a Liquid Chromatography-Tandem Mass Spectrometry (LC-MS/MS) analysis. The digest was measured by means of a Q-Exactive™ Focus Hybrid Orbitrap mass spectrometer equipped with a Thermo Scientific™ Vanquish™ ultra-high performance liquid chromatography system (Thermo Fisher Scientific). 5 µL of the tryptic digests were injected and chromatographically separated by means of 35 min linear gradient of 2-45% mobile phase B (mobile phase A: 0.1% formic acid, mobile phase B: 0.1% formic acid in acetonitrile) on an Acquity UPLC® CSH C18 column (2.1 × 150 mm, 1.7 µm) from Waters (Milford, MA, USA). The flow rate and column temperature were 0,3 mL/min and 45 °C, respectively. The Q-Exactive Focus, operating in data dependent acquisition mode (DDA), was set to perform a mass spectrometry (MS) scan (R= 70 000 at 200 m/z, AGC target 3.0 e6) from 375 to 1500 m/z, followed by HCD MS^2^ spectra (R= 17 500 at 200 m/z, NCE =27%, AGC target = 1.0e5, max ion time = 50 ms) on the three most abundant precursors (quadrupole isolation width 1.4 m/z). Data treatment and data analysis were performed by means of PEAKS Studio 11 (Bioinformatics Solutions Inc., Waterloo, Canada). *De novo* sequencing and database searches were performed with a 10-ppm precursor mass tolerance, a 0.02 Da fragment tolerance and an FDR <0.1% on the peptide level. Oxidations of methionine were set as a variable modification while the carbamidomethylation of cysteines was included as a fixed modification. Only fully tryptic peptides and a maximum of three trypsin miscleavages were allowed.

### Analytical Size Exclusion Chromatography

Analytical Size Exclusion Chromatography (SEC) was performed using a Superdex® 200 Increase 10/300 GL column (Cytiva) equilibrated with 10 mM CAPS, pH 11.0. To calibrate the column and estimate the molecular weight of the eluted species, a Gel Filtration Standard Mix (#1511901, Bio-Rad) was used. This standard mixture includes thyroglobulin (670 kDa), bovine γ-globulin (158 kDa), chicken ovalbumin (44 kDa), equine myoglobin (17 kDa), and vitamin B12 (1.35 kDa). A standard calibration curve was generated by plotting the distribution coefficient (K_av_) of each standard protein against the logarithm of its molecular weight, allowing estimation of the apparent molecular weight of hnRNPA2B1.

### Mass Photometry

Protein landing was recorded using a Refeyn OneMP (Refeyn Ltd) MP system by diluting 10 μL of a 150 nM hnRNPA2B1 solution into a 10 μL drop of filtered 10 mM CAPS pH 11.0 buffer (Final concentration: 75 nM). Movies (6,000 frames, 60 s) were acquired with the AcquireMP software version 2.1.1 (Refeyn Ltd) using the default settings. Data was analysed using default settings on DiscoverMP (version 2.1.1; Refeyn Ltd). Contrast-to-mass calibration was performed with MassFerence P1 (Refeyn) using standards of 88, 172, 258, and 344 kDa. The binding and unbinding events were grouped into mass ranges (binning) using GraphPad Prism. Frequency distribution with default setting and a bin width of log2(x), where x is the total number of detected particles, was used. Data was represented as the number of particles (counts) vs. mass (kDa). Triplicates were measured, and the average molecular weight and standard deviation were calculated.

### Turbidity measurements

hnRNPA2B1 remains soluble in 10 mM CAPS buffer at pH 11.0. LLPS was induced by diluting the stock solution with 100 mM HEPES buffer (pH 7.5) to a final protein concentration of 10 µM. After mixing, the solutions were pipetted into a black, non-binding, flat-bottom 384-well microplate (Greiner Bio-One) and sealed with VIEWseal™ adhesive film (Greiner Bio-One). Turbidity was recorded every 4 minutes at 600 nm and 340 nm using a Synergy™ HTZ Multi-Mode Microplate Reader (BioTek). Measurements were performed at 25 ºC with slow, continuous agitation for 2 hours. All experiments were conducted with at least three technical replicates. Baseline correction was applied by subtracting buffer-only controls.

### Dynamic Light Scattering

Dynamic Light Scattering (DLS) measurements were performed using a DynaPro® NanoStar® instrument (Wyatt Technology). After inducing LLPS at the same conditions as in the turbidity measurements, a disposable cuvette (Wyatt) was filled with 100 µL of phase-separating hnRNPA2B1, and the outer well was filled with water. The cuvette was sealed to prevent evaporation. Scattered light intensity was recorded at a fixed angle (95º) with 5 acquisitions of 8 seconds each. Measurements were collected over 2 hours at 25ºC. Data were analyzed using the Dynamics® Software Package (V. 7.10.1.21, Wyatt). The acquisitions were used to calculate the hydrodynamic radius based on the autocorrelation function, ultimately corresponding to a single time point.

The translational diffusion coefficient (D_t_) was calculated using the Stokes-Einstein equation:

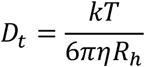

Where *k* is the Boltzmann constant, *T* is the temperature, *η* is the viscosity (assumed to be that of water), and *R*_*h*_ is the measured hydrodynamic radius. All experiments were performed in triplicate.

### Brightfield and Fluorescence Microscopy

Microscopy measurements were performed on a Leica DMi8 inverted microscope equipped with a Leica DFC7000 GT camera. Protein solutions were supplemented with 25 µM thioflavin T (ThT) as a fluorescent dye. Phase separation was induced, and hnRNPA2B1 droplets were visualized with 100X oil-immersion objectives under both brightfield and fluorescence modes (FITC filter). Images were captured with Leica Application Suite X (LAS X) software (Leica Microsystems) and analyzed with Fiji (ImageJ).

### Data visualization and analysis

Plasmid maps were created with SnapGene version 8.1. Raw data were preprocessed in Microsoft Excel 2024 before graphing and statistical analysis in GraphPad Prism version 10.5.0.

## 3. Results and Discussion

### Sequence-based protein disorder, aggregation, and LLPS propensity prediction for hnRNPA2B1

To guide our experimental design and gain region-based insights into the structural properties of hnRNPA2B1, we performed sequence-based bioinformatic predictions [**Fig. 1**]. Intrinsic disorder prediction using the latest-generation AIUPred tool [20] confirms the literature-derived information on hnRNPA2B1 being extensively disordered [**Fig. 1A**], particularly within its C-terminal LCD. Only the first RRM is predicted to form a stable, globular fold, while the second RRM shows a lower propensity for stable folding [**Fig. 1A**].

**Figure 1:**
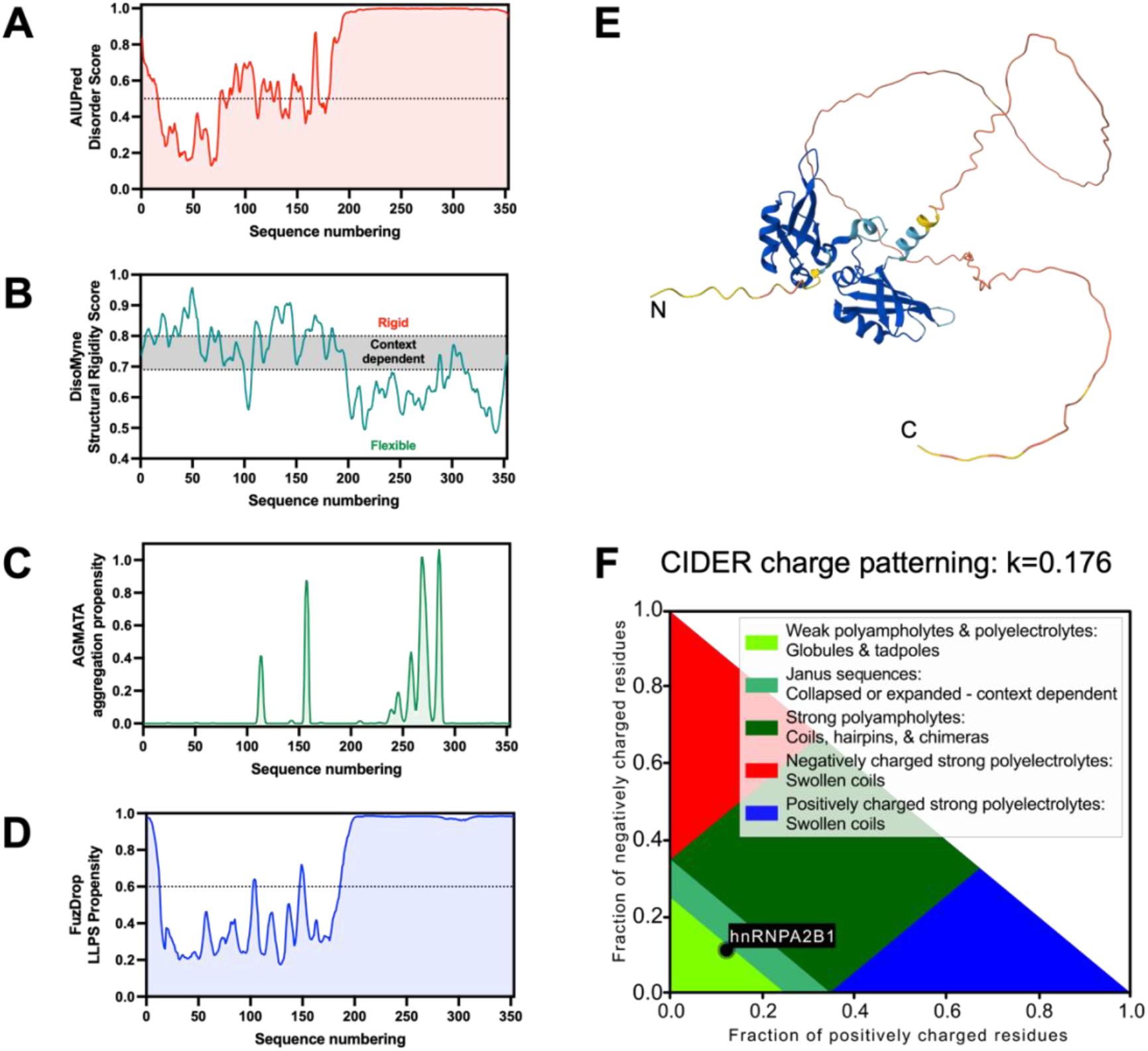
hnRNPA2B1 is predicted to be partially disordered with a C-terminal low complexity domain of moderately high aggregation propensity and very high liquid-liquid phase separation propensity. (A) AIUPred structural disorder prediction; (B) DynaMine backbone S^2^ order parameter prediction for structural rigidity or flexibility; (C) Prediction of aggregation-prone regions by AGMATA (plotted with a scaling factor of 0.3 – default value); (D) Prediction of liquid-liquid phase separation-prone regions by FuzDrop; (E) AlphaFold2 model of hnRNPA2 colored by the pLDDT model confidence score; and (F) CIDER-based state diagram for the predicted collapse of the polypeptide chain and the calculated kappa (κ) charge patterning parameter.

DynaMine [21,22] predictions further support this domain organization, indicating that the N-terminal half of the protein is more rigid (although mostly dependent on the context) than the highly flexible LCD [**Fig. 1B**]. Aggregation hotspot predictions using AGMATA [24] reveal that most aggregation-prone regions are located within the LCD [**Fig. 1C**], aligning with its known role in amyloid-like assembly formation. Furthermore, the FuzDrop algorithm [25] predicts that the entire LCD is highly LLPS prone [**Fig. 1D**], thus it can drive and scaffold the homotypic phase separation of the protein.

Structural modeling using the AlphaFold2-predicted tertiary structure of hnRNPA2B1 [26] shows high confidence (pLDDT > 90, dark blue) for the RRMs, which are well-folded into stable structures, whereas the LCD and linker regions display low to very low model confidence (pLDDT < 70, yellow and orange) [**Fig. 1E**], consistent with intrinsic disorder [29–31]. Interestingly, although the AlphaFold2 model does not suggest a high level of structural collapse mediated by contacts between distant regions, CIDER [23] analysis suggests a higher level of compactness for the protein. Based on the distribution of positively and negatively charged residues, hnRNPA2B1 is predicted to adopt a partially disordered, somewhat globular shape or context-dependent collapsed disordered chain [**Fig. 1F**]. However, due to its moderately low charge segregation, its kappa value (κ*=*0.176) can be interpreted as a partially collapsed molten globule with context dependence [23].

For example, LLPS would certainly qualify as a context where a higher level of compactness may appear, as coil-to-globule transition has been associated with liquid-liquid phase separation [32,33]. Together, these sequence-based predictions highlight the extensive disorder and strong propensity for phase separation and aggregation of hnRNPA2B1, which pose challenges for recombinant expression and purification.

### Expression and Purification Protocol for hnRNPA2B1

To overcome solubility and degradation issues associated with the LCD, we employed C-terminal fusion of a SUMO tag, which increases solubility, protects disordered regions, and enhances protein expression [34]. The SUMO-hnRNPA2B1 fusion was cloned into the pET-22b (+) vector, incorporating a C-terminal hexahistidine tag for affinity purification **[Fig. 2A, Fig. S1]**. Robust protein expression is achieved following induction with 1 mM IPTG for 4 hours at 37ºC. SDS-PAGE analysis of total cell lysates revealed a prominent band around 55 kDa, consistent with the expected size of the fusion protein (theoretical MW: 50.7 kDa) **[Fig. 2B]**. The slower migration and higher apparent MW compared to the theoretical size are likely due to the protein’s intrinsically disordered regions (IDRs), which bind SDS less efficiently and thus migrate more slowly than globular proteins of similar mass [34]. Western blotting using an anti-polyHistidine antibody confirms the identity of the band as SUMO-hnRNPA2B1 **[Fig. 2C]**.

**Figure 2:**
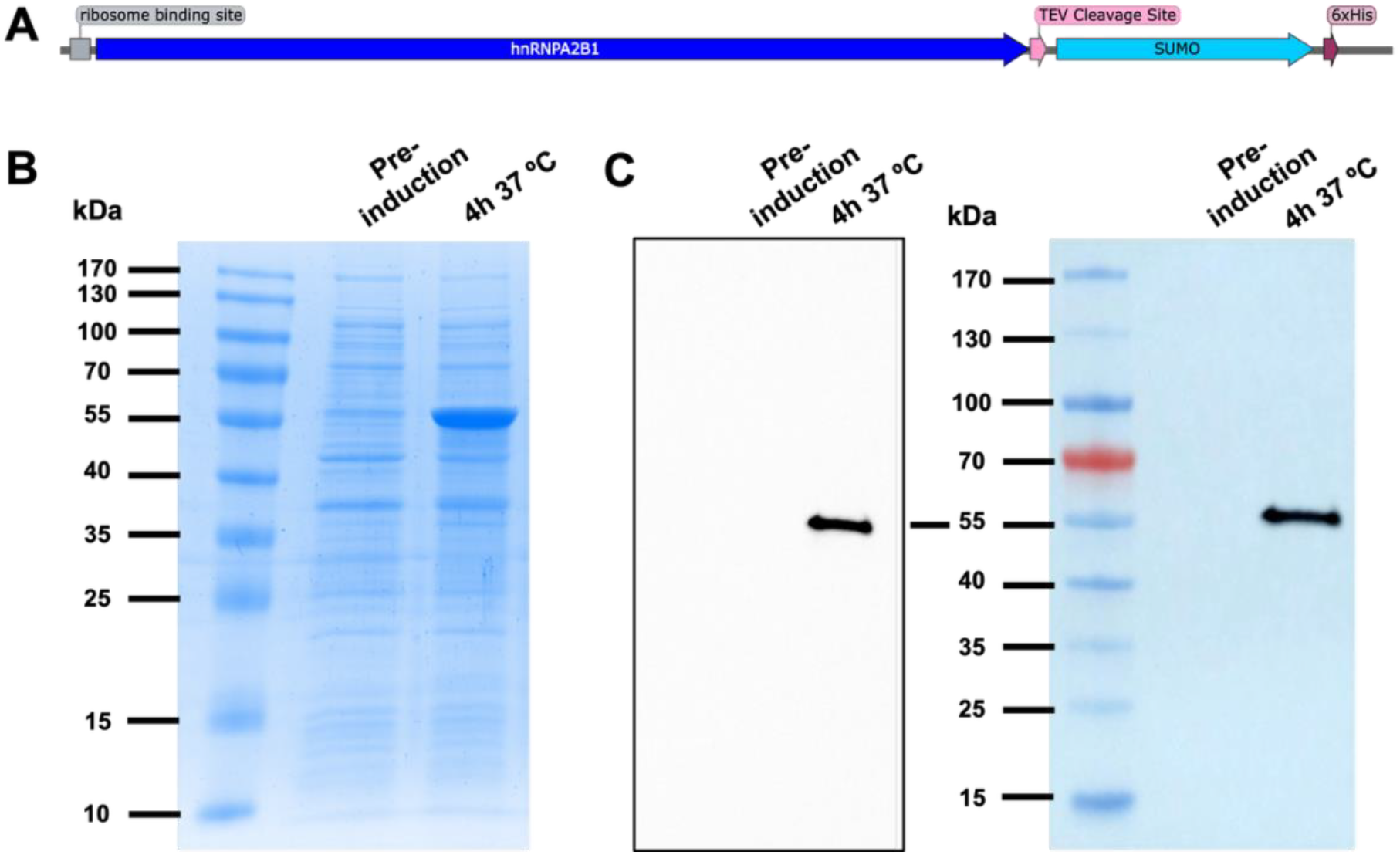
Expression of recombinant SUMO-hnRNPA2B1. (A) Schematic representation of the ORF in the pET-22b (+) expression vector used for bacterial production of SUMO-hnRNPA2B1. (B) SDS-PAGE of total cell lysates after 4-hour induction with 1 mM IPTG at 37 °C reveals a prominent band at ∼55 kDa, consistent with the predicted size of SUMO-hnRNPA2B1 (theoretical MW: 50.7 kDa). (C) Western blot using an anti-His antibody confirms the 55 kDa band as polyHistidine-tagged SUMO-hnRNPA2B1, with no detectable nonspecific binding. The first image is the chemiluminescent revealing of the membrane. The second image overlaps the chemiluminescence and the protein ladder from the same membrane.

SUMO-hnRNPA2B1 was purified from the pellet of a 1 L *E. coli* culture. Given that hnRNPA2B1 is an RNA-binding protein, minimizing nucleic acid contamination is essential to avoid artifacts in downstream experiments. Therefore, the lysis buffer was supplemented with DNAse I and RNAse A, and a high salt concentration (1M NaCl) was used to disrupt electrostatic interactions with nucleic acids. Following sonication, the cellular debris was removed by centrifugation, and the clarified lysate containing SUMO-hnRNPA2B1 was filtered and loaded onto a nickel-charged column **[Fig. 3, Fig. S2]**. Eluted fractions were desalted by buffer exchange into a cleavage buffer, supplemented with TEV protease, and incubated overnight. After tag removal, hnRNPA2B1 remains soluble due to the high ionic strength of the buffer, which disrupts electrostatic interactions between the charged residues and prevents protein aggregation. The cleaved protein was separated from the protease and other contaminants using a reverse nickel-affinity purification step, collecting the flowthrough containing tag-free hnRNPA2B1 **[Fig. 3]**. Protein concentration and purity were assessed by UV absorbance after each chromatographic step using the corresponding extinction coefficients. After the final dialysis step, approximately 3 mg of pure hnRNPA2B1 were obtained per liter of culture, with minimal nucleic acid contamination (A_260_/A_280_ < 0.6) [**Table S1]**.

**Figure 3:**
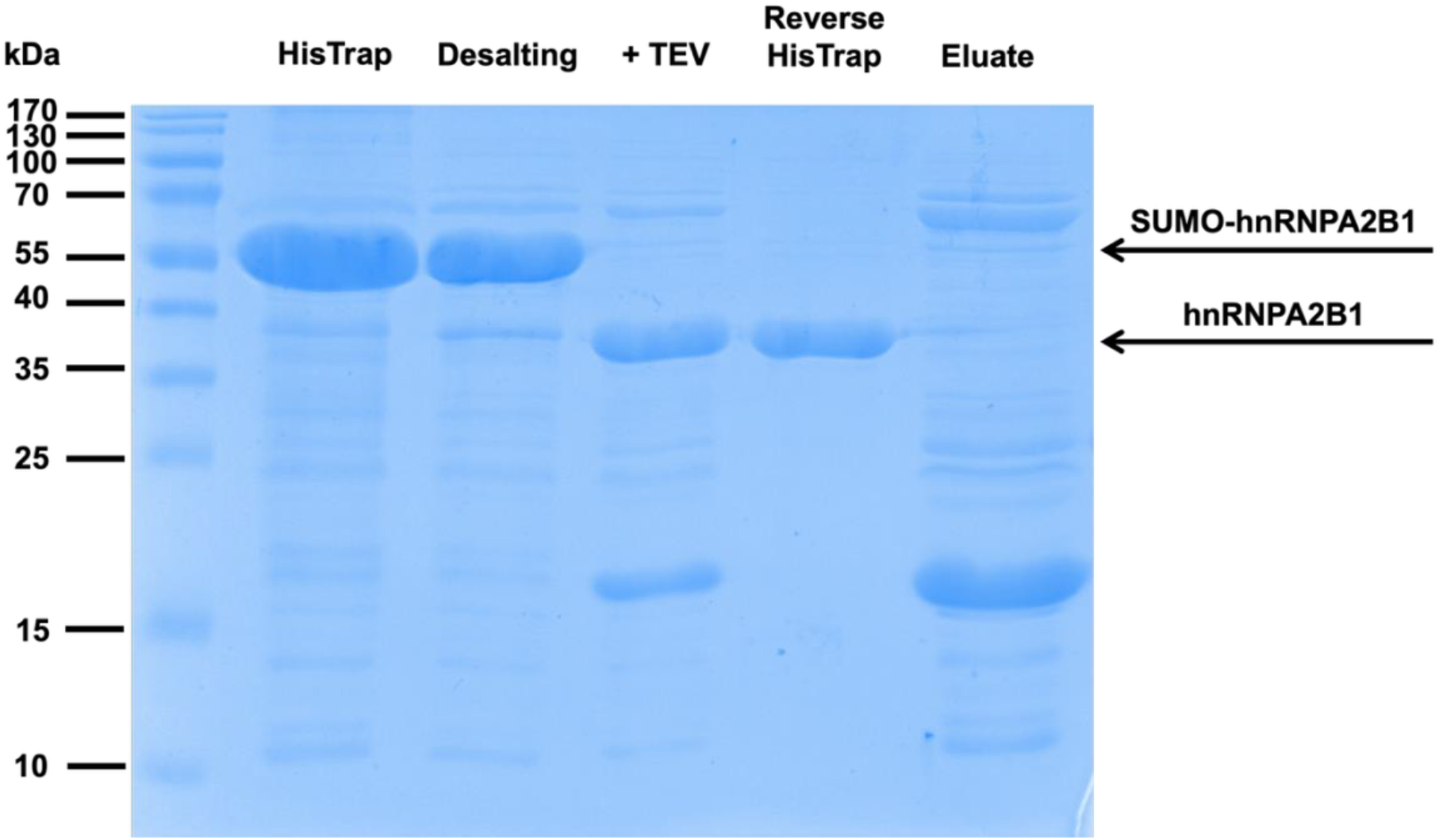
Purification of full-length hnRNPA2B1. SDS PAGE of protein samples after each chromatographic step. Prominent bands in the first HisTrap and desalting steps correspond to SUMO-hnRNPA2B1 (theoretical MW: 50.7 kDa). Upon addition of TEV protease, the SUMO tag is efficiently cleaved. After a second HisTrap, the flow-through contains pure, cleaved, and soluble hnRNPA2B1 (theoretical MW: 38.2 kDa). Remaining contaminants are present in the eluate.

To identify a suitable storage buffer, we analyzed the net charge-pH profile of hnRNPA2B1 **[Fig. 4A]**. At high pH, hnRNPA2B1 adopts a strong negative net charge due to deprotonation of acidic and some basic residues, which inhibits condensation. In contrast, at near-native pH (∼7.0-7.5), an intermediary net charge is expected to promote LLPS **[Fig. 4B]**. This “pH jump” strategy allows LLPS-prone proteins to be stored in solution at high pH and induced for LLPS under near-native conditions without denaturants, solubility tags, or high salt [14]. It is important to note that at high pH, the hydroxyl groups of tyrosine residues (pKa ∼10.9) become deprotonated, which can artificially elevate the A_260_ signal and skew the A_260_/A_280_ ratio [35]. Therefore, nucleic acid contamination was assessed before the final dialysis step.

**Figure 4:**
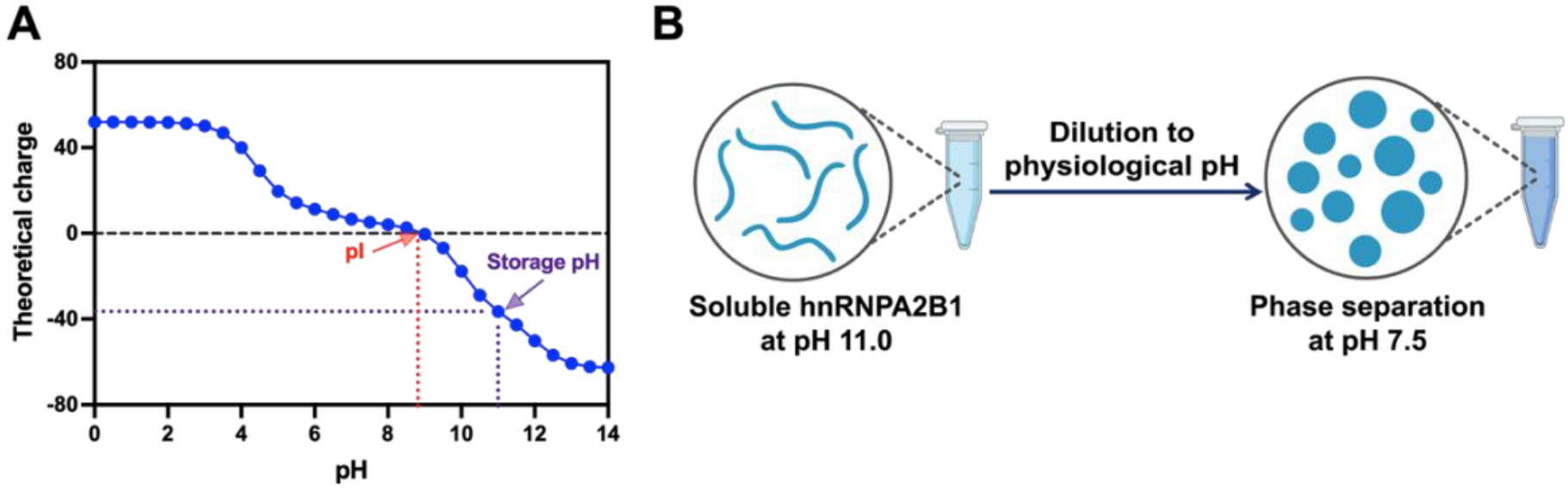
pH modulating the solubility and phase separation of hnRNPA2B1. (A) The theoretical net charge of full-length hnRNPA2B1 was calculated using the ProtCalc web server. hnRNPA2B1 has a theoretical isoelectric point of 8.8 (red line). At basic pH, acidic residues, tyrosine, and basic residues like lysine become deprotonated, increasing the negative charge of the protein. (B) Schematic representation of the pH jump method to induce LLPS at near-native conditions. LLPS can be easily initiated by diluting the high pH hnRNPA2B1 solution into a buffer at physiological pH (7.0-7.5).

### Confirmation of purified hnRNPA2B1

To confirm the identity and the purity of the purified protein, we performed both MALDI-TOF and LC-MS/MS experiments. Firstly, intact mass measurements, acquired between 10 000 and 100 000 m/z, yielded a mass spectrum for which all main peaks corresponded to different charge states of intact hnRNPA2B1. Moreover, no other significant peaks were detected that did not correspond to hnRNPA2B1. External calibration of more accurate acquisitions yields a measured mass of 38223 ± 4 Da **[Fig. 5]**, in agreement with the expected molecular weight of hnRNPA2B1 following cleavage of the solubility tag (theoretical MW: 38225 Da). Secondly, both top-down sequencing (TDS) by means of MALDI-ISD-TOF and an LC-MS/MS analysis of a trypsin digest of the purified protein were performed to verify the primary structure. Both experiments confirm the amino acid sequence. More concretely, the ion-series obtained in the MALDI-ISD-TOF MS spectrum match both theoretical fragments (c-, y- and z+2-ions) of the N- and C-terminus of hnRNPA2B1 **[Fig. S3A, B]**, while up to 72% of the entire amino acid sequence was identified on the peptide-level by LC-MS/MS analysis **[Fig. S3C]**. Combined, these experiments indicate that hnRNPA2B1 is intact, of high purity and validate the effectiveness of our purification strategy and confirm that the protein is suitable for downstream assays.

**Figure 5:**
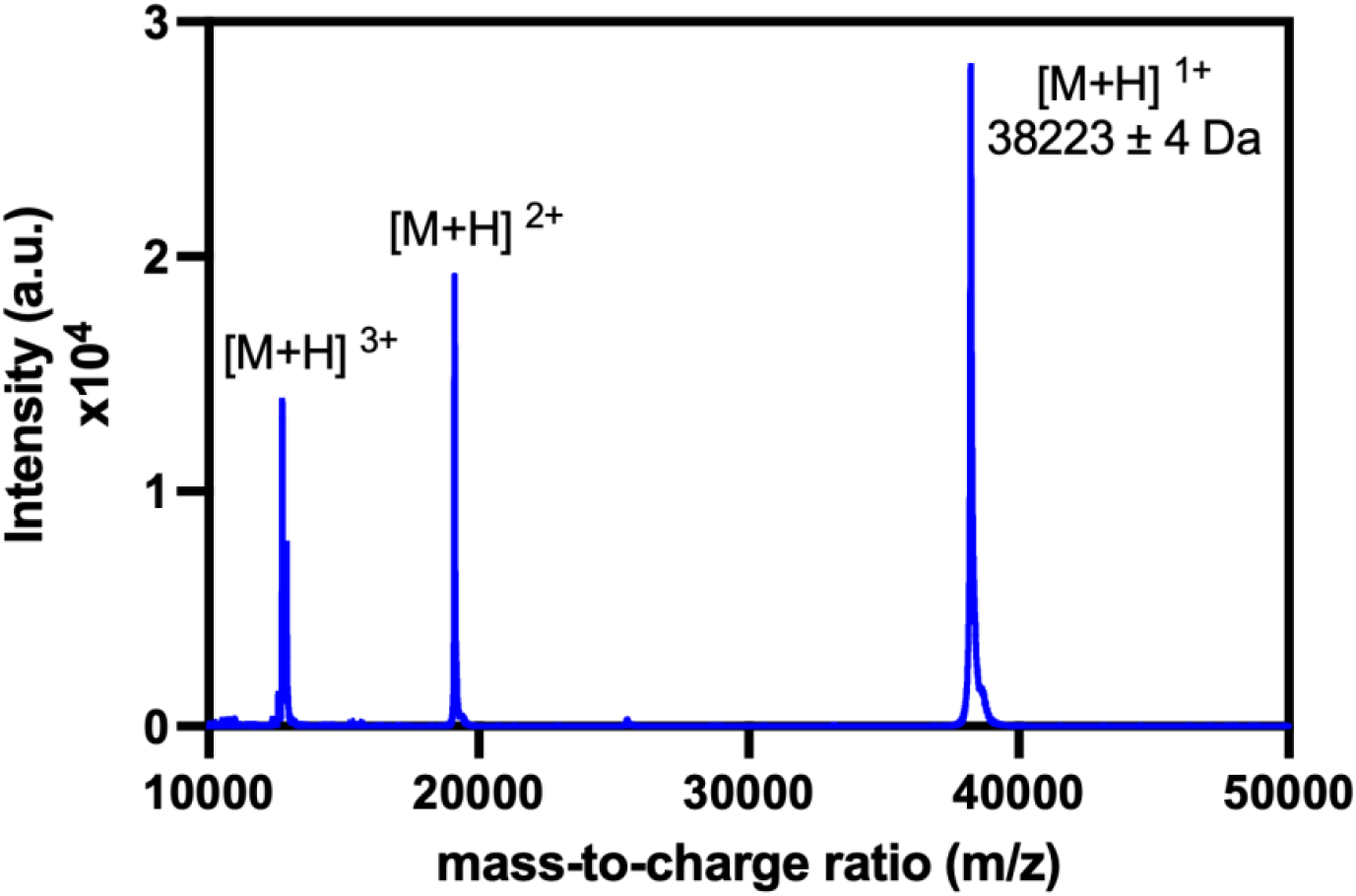
MALDI-TOF mass spectrum of intact hnRNPA2B1. The spectrum shows a dominant peak at m/z ≈ 38223, corresponding to the single-charged molecular ion [M+H]^1+^ of full-length hnRNPA2B1. This mass is consistent with the theoretical MW of hnRNPA2B1 after tag cleavage (MW: 38,225 Da), confirming the identity and the purity of the protein.

We further assessed the oligomeric state of purified hnRNPA2B1 in the high-pH storage buffer by combining Mass Photometry with Analytical Size Exclusion Chromatography (SEC). Mass Photometry measurements reveal a predominantly dimeric species (*∼* 78 kDa) and a much smaller population of a tetrameric state (*∼*153 kDa) **[Fig. 6A]**. We complemented this assay with Analytical Size Exclusion Chromatography (SEC) using a 15 µM hnRNPA2B1 solution. The SEC profile reveals a main peak at 11 mL **[Fig. 6B]**, which corresponds to an apparent molecular weight of *∼* 93 kDa when compared to the calibration curve **[Fig. S4]**. This peak has a preceding shoulder around 10 mL, which likely represents a trimer or a tetramer (*∼* 130 kDa). The discrepancy between the calculated molecular weights from SEC and mass photometry, may be a consequence of the intrinsically disordered regions in hnRNPA2B1. IDRs can lead to early elution in SEC, altering the apparent molecular weight. These results ultimately suggest that at pH 11.0, hnRNPA2B1 exists as a soluble protein predominantly in oligomeric forms. Whereas the low-complexity domain of hnRNPA2B1 mediates self-association in the context of LLPS and aggregation, dimerization of the RNA Recognition Motifs has been reported in the presence of nucleic acids [36,37]. It is possible that the high pH of the storage buffer may also promote weak, transient oligomerization of hnRNPA2B1 in absence of nucleic acids. Despite the oligomeric state of hnRNPA2B1 at high pH, the protein remains soluble and suitable for downstream applications.

**Figure 6:**
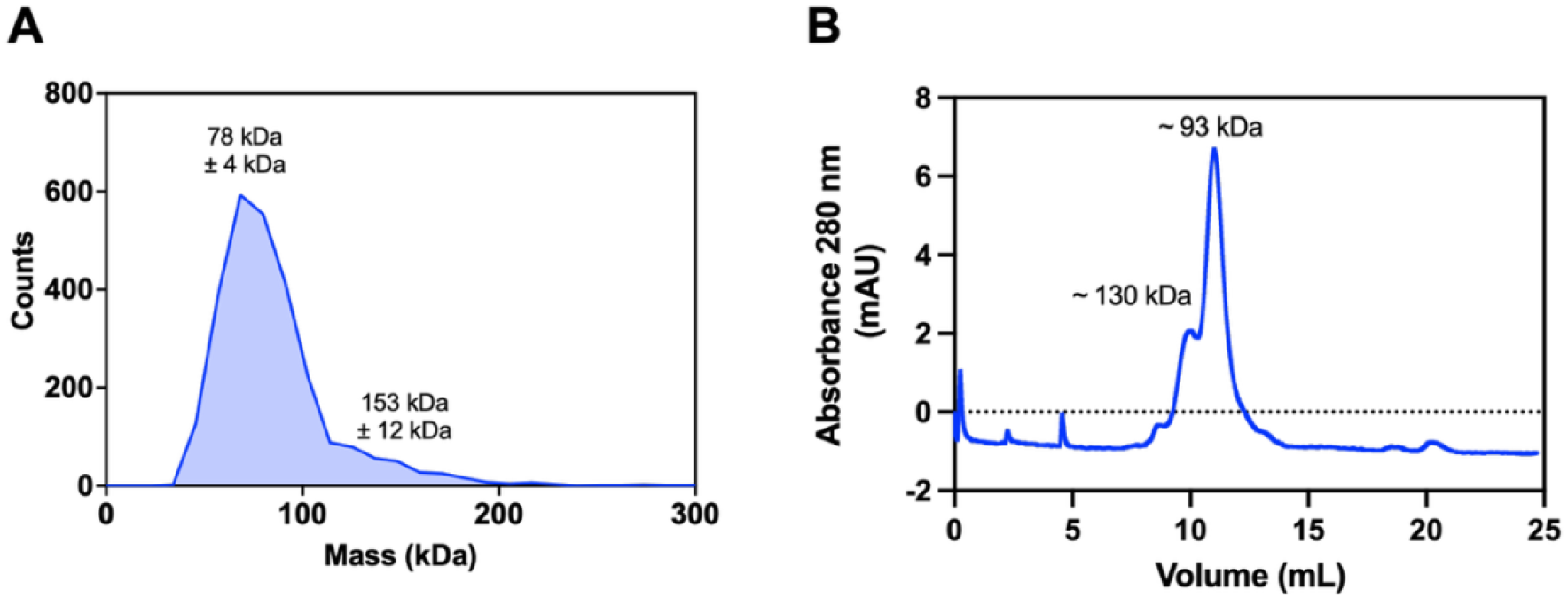
Oligomeric state of hnRNPA2B1 in the storage buffer. (A) Mass Photometry data shows that hnRNPA2B1 exists as a dimer (∼ 78 kDa) with a smaller tetrameric population (∼ 153 kDa) in high-pH storage buffer. (B) Analytical Size Exclusion Chromatography of hnRNPA2B1. The predominant peak has a small shoulder corresponding to a higher-order oligomer, potentially the tetrameter, before the main population, likely the dimer is detected.

### hnRNPA2B1 undergoes LLPS and forms liquid-like droplets

A primary functional test of hnRNPA2B1 is to demonstrate its phase separation propensity. We induced LLPS using the pH jump method [14], by diluting the protein stock to 10 µM in 0.1 M HEPES buffer at pH 7.5. Upon dilution, the solution transitioned from clear to turbid. The kinetic path of LLPS was followed by measuring the turbidity of the solution at two different wavelengths: 340 nm and 600 nm. At both wavelengths, a rapid increase in turbidity is observed with a maximum reached in minutes, followed by a slower decrease before reaching equilibrium **[Fig. 7A]**. This profile is consistent with droplet maturation and sedimentation. Centrifugation of the phase-separated solution and further SDS-PAGE showed hnRNPA2B1 to be localized in the pellet, with no detectable protein remaining in the supernatant **[Fig. 7B]**, confirming its incorporation into the droplets.

**Figure 7:**
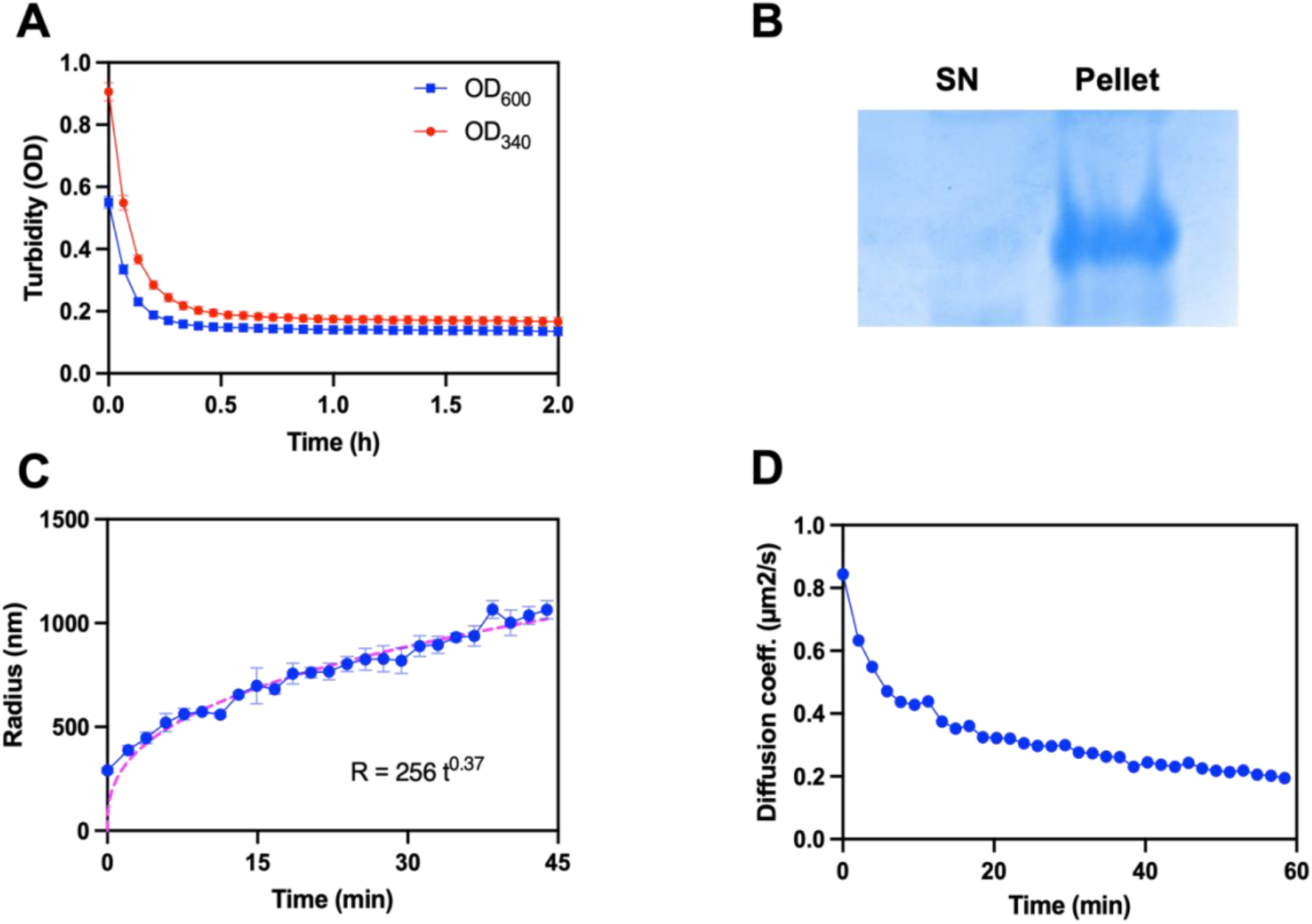
Kinetic trace of liquid-liquid phase separation of hnRNPA2B1. LLPS of 10 µM hnRNPA2B1 solutions was induced by a pH jump from 11.0 to 7.5 and followed through different techniques. (A) Turbidity measurements at 340 nm (red line) and 600 nm (blue line) show that the maximum in absorbance is reached minutes after inducing LLPS, followed by a gradual decrease until equilibrium is reached. (B) SDS-PAGE of hnRNPA2B1 samples showing that the protein localizes to the pelleted droplets, leaving the supernatant devoid of detectable protein. (C) hnRNPA2B1 droplets grow as a function of time (t^1/3^), following Ostwald ripening, in which larger droplets grow at the expense of the smaller ones. After one hour, the data could not be interpreted, probably due to protein aggregation. (D) The decrease in the diffusion coefficient of hnRNPA2B1 droplets shows that, over time, they transition from a liquid-like behavior to a more gel-like or solid-like state.

To measure directly the time course of droplet growth and maturation, we performed dynamic light scattering (DLS) experiments. DLS relates the Brownian motion of macromolecules in solution to particle size, allowing us to monitor the changes in droplet size over the LLPS kinetics. Immediately after inducing LLPS, small droplets with a hydrodynamic radius of ∼ 300 nm form. Over the next 30 to 45 minutes, droplet size increases following a power-law exponent of ⅓, characteristic of Ostwald ripening **[Fig. 7C]**. In Ostwald ripening, the larger droplets grow at the expense of the smaller ones [38,39]. After approximately one hour, the droplets reached a radius of ∼ 1500 nm. Beyond this point, the DLS data become uninterpretable due to increased polydispersity, likely caused by droplet fusion, transition to aggregates, and precipitation in the cuvette **[Fig. S5]**.

Analysis of the translational diffusion coefficient (*D*_*t*_) over time further supports this transition towards aggregation. Initially, the first droplets exhibit a *D*_*t*_ (∼0.84 µm^2^/s) consistent with liquid-like behavior. As droplets grow and mature, *D*_*t*_ decreases, reflecting both increased size [40] and a transition toward gel-like or solid-like states, where internal protein mobility is reduced [41] **[Fig. 7D]**.

### hnRNPA2B1 droplets mature into amorphous aggregates

To directly visualize droplet formation, growth, and subsequent aggregation of hnRNPA2B1, we performed fluorescence microscopy using Thioflavin T (ThT) as a fluorescent probe. ThT is a well-established dye for detecting β-sheet-rich structures, including both phase-separated droplets [10,42] and protein aggregates [5,11,18], and its green fluorescence emission (∼482 nm) [43] allows imaging with standard FITC/GFP filter sets.

Following the induction of LLPS, we observed the rapid formation of small, spherical, and highly mobile, liquid-like droplets **[Fig. 8A, B]**. The droplets bind ThT due to potential early structural ordering into β-sheet structures, a process mediated by the low-complexity domain, which contains different motifs that mediate self-association and fibrillization [5,6,9,12] Over time, the droplets gradually undergo a morphological transition into irregular, ThT-positive aggregates. While ThT is mostly commonly used to detect amyloid-like fibrils, the aggregates formed by hnRNPA2B1 likely contain sufficient β-sheet structures to support strong ThT binding. A loss of sphericity is evident approximately 30 minutes post-LLPS [**Fig. 8C]**, coinciding with the appearance of amorphous aggregates, indicative of an irreversible transition to a solid-like or gel-like state. By two hours post-LLPS, the field of view is dominated by amorphous aggregates **[Fig. 8D, E]** with no remaining spherical droplets.

**Figure 8:**
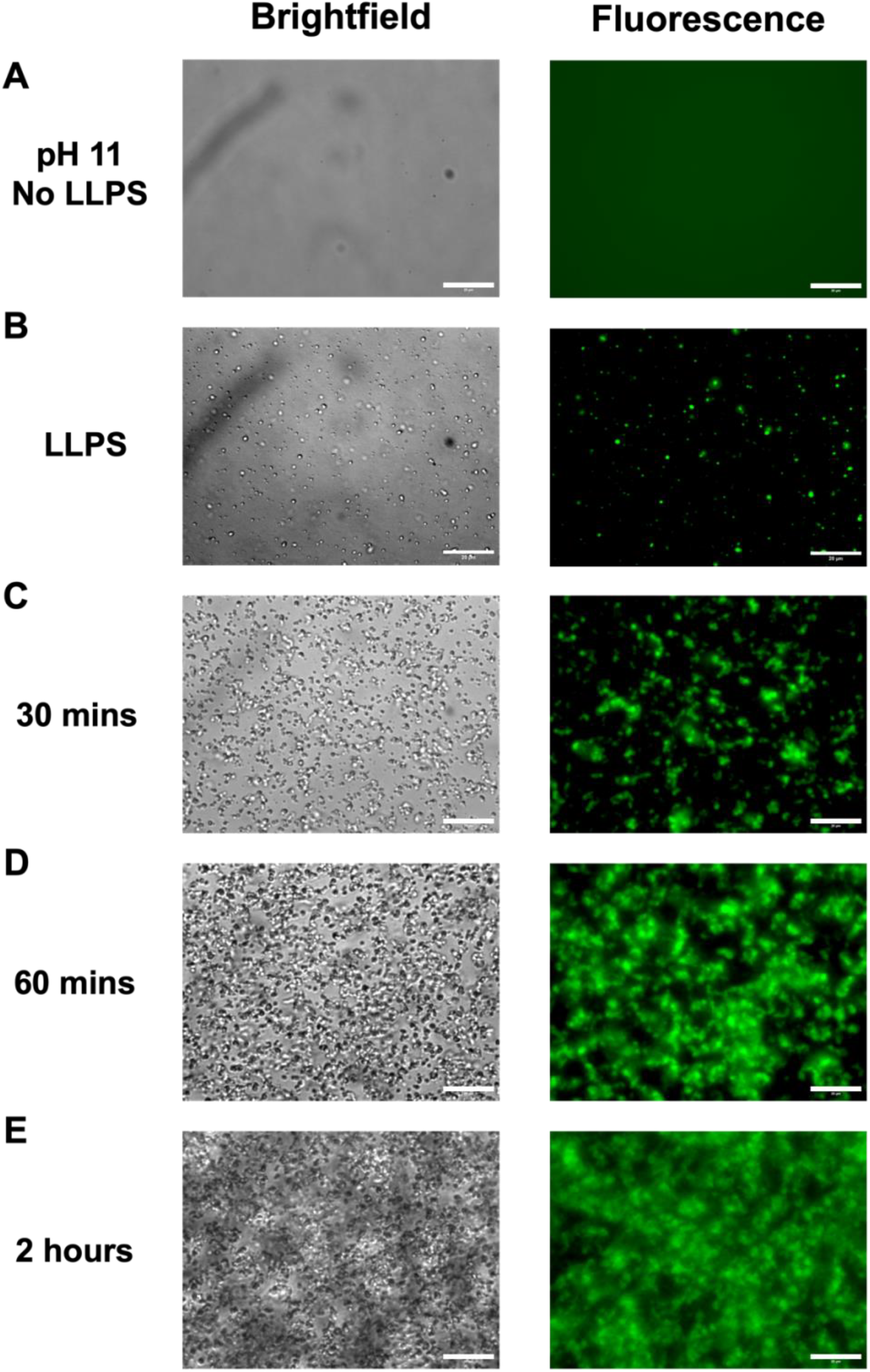
hnRNPA2B1 droplets transition into amorphous aggregates. LLPS of 10 µM hnRNPA2B1 supplemented with 25 µM Thioflavin T was visualized with Brightfield and Fluorescence microscopy. (A) At pH 11.0, hnRNPA2B1 remains in solution, with no visible droplets or aggregates. (B) Upon inducing LLPS, spheric, dynamic, liquid-like droplets formed. (C) After 30 minutes, the droplets began losing their sphericity and the first amorphous aggregates appeared. After one (D) and two (E) hours, all droplets had become irreversible, amorphous aggregates. Scale bar: 20 µM.

The timescale of this liquid-to-solid transition is consistent with our turbidity and DLS measurements, which indicate that hnRNPA2B1 droplets remain liquid-like for around 30 minutes following LLPS induction. The observed droplet growth and transition into amorphous aggregates align with the increase in polydispersity and the decrease in diffusion coefficient over time. Together, these findings support a model in which hnRNPA2B1 undergoes a maturation process from dynamic condensates to amorphous aggregates, reinforcing previous the hypothesis that LLPS precedes protein aggregation [42,44].

## 4. Conclusions

In this study, we developed a robust protocol for the expression and purification of full-length hnRNPA2B1, an RNA-binding protein with an aggregation-prone, intrinsically disordered region. By combining a high-salt purification strategy with an extreme-pH storage buffer, we effectively remove nucleic acid contamination and maintain the protein in solution even after cleavage of the solubility tag. This protocol yields protein of sufficient purity for downstream biophysical characterization of LLPS, a process highly sensitive to experimental conditions.

Our *in vitro* characterization revealed that purified hnRNPA2B1 undergoes LLPS upon a pH jump, forming dynamic, ThT-positive droplets that progressively transition into amorphous aggregates. This maturation process was consistently observed across multiple techniques, supporting a model in which LLPS serves as a precursor to protein aggregation.

The protocol presented here will facilitate further studies on hnRNPA2B1 in the context of LLPS and aggregation, potentially providing mechanistic insights into the transition from functional condensates into pathogenic assemblies. Beyond hnRNPA2B1, this workflow may be applicable to purify other intrinsically disordered, aggregation-prone proteins, without the need for denaturants or persistent solubility tags, enabling downstream biophysical and functional studies under native-like conditions.

## CRediT authorship contributions

**Luis Fernando Durán-Armenta:** Investigation, methodology, formal analysis, visualization, writing - original draft, writing - review and editing. **Attila Mészaros:** Investigation, methodology, formal analysis, writing - review and editing. **Julia Malo Pueyo:** Investigation, formal analysis, writing - review and editing. **Steven Janvier:** Investigation, formal analysis, writing - review and editing. **Tamas Lazar:** Software, Formal analysis, visualization, writing - original draft, writing - review and editing. **Dominique Maes:** Conceptualization, supervision, writing - review and editing. **Remy Loris:** Conceptualization, supervision, writing - review and editing. **Peter Tompa:** Conceptualization, supervision, writing - original draft, writing - review and editing.

## Declaration of generative AI and AI-assisted technologies in the writing process

During the preparation of this work, the author(s) used Microsoft Copilot to assist with language editing, grammar correction, and improving the clarity and flow of the manuscript text. After using this tool/service, the author(s) reviewed and edited the content as needed and take(s) full responsibility for the content of the published article.

## Data availability

Data will be made available upon reasonable request.

## Acknowledgments

The authors thank Joris Van Lindt (VIB-KU Leuven Center for Brain Disease) for his helpful insights and suggestions during the initial stages of the project, and Sarah Haesaerts (VIB-VUB Center for Structural Biology and Structural Biology Brussels, Vrije Universiteit Brussel) for her assistance with the Analytical Size Exclusion Chromatography.

