## Supplementary Material for "Expression and Purification of Full-Length hnRNPA2B1 for Biophysical Characterization of Liquid-Liquid Phase Separation"

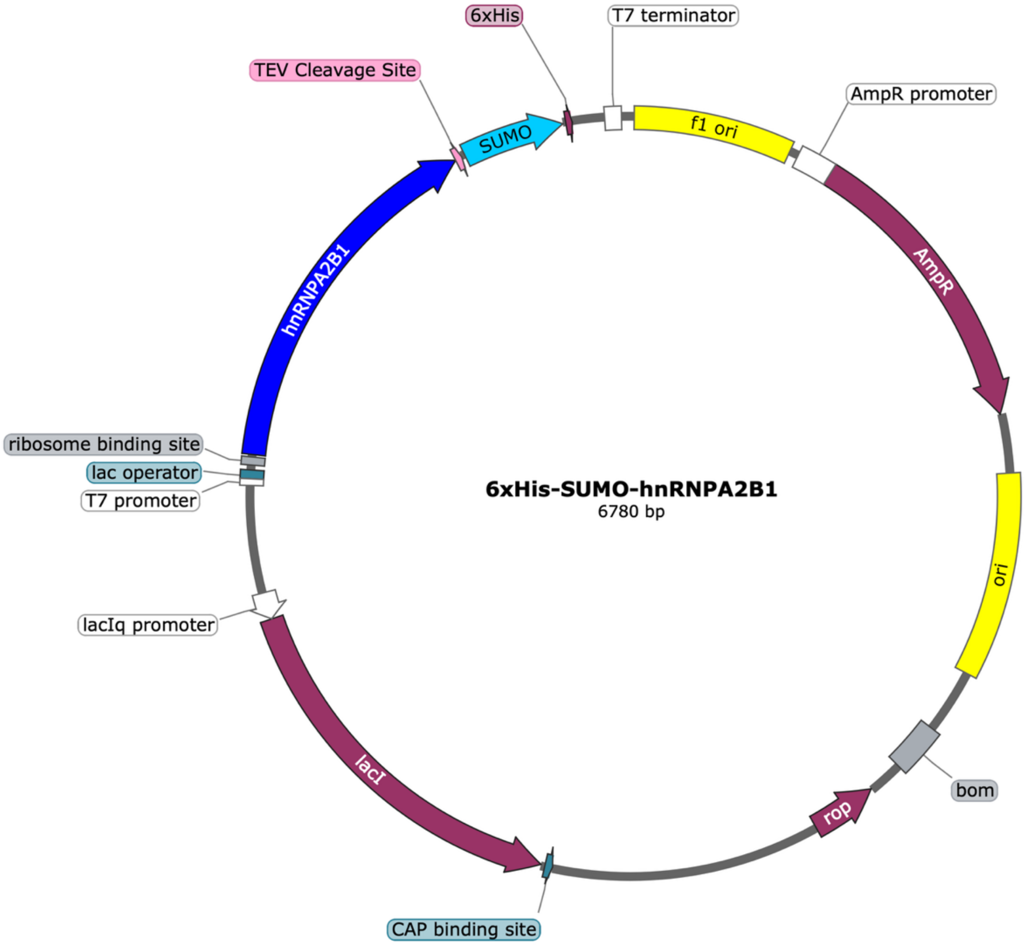


**Figure S.1. Plasmid map of the pET22B(+) encoding SUMO-hnRNPA2B1 used for the recombinant expression of hnRNPA2B1 in *E. coli.***

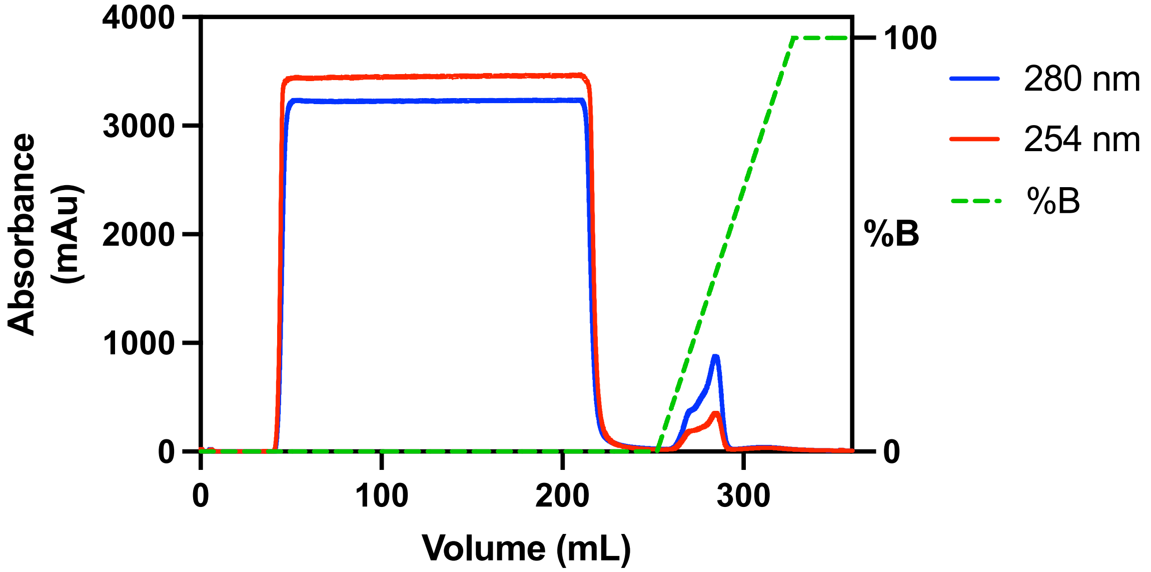


**Figure S.2. Ni.NTA chromatography of hnRNPA2B1.**
*The UV absorbance at 280 nm (blue) and 260 nm (red) is indicated. The right axis and green line indicate the percentage of high imidazole buffer used to elute the protein.*

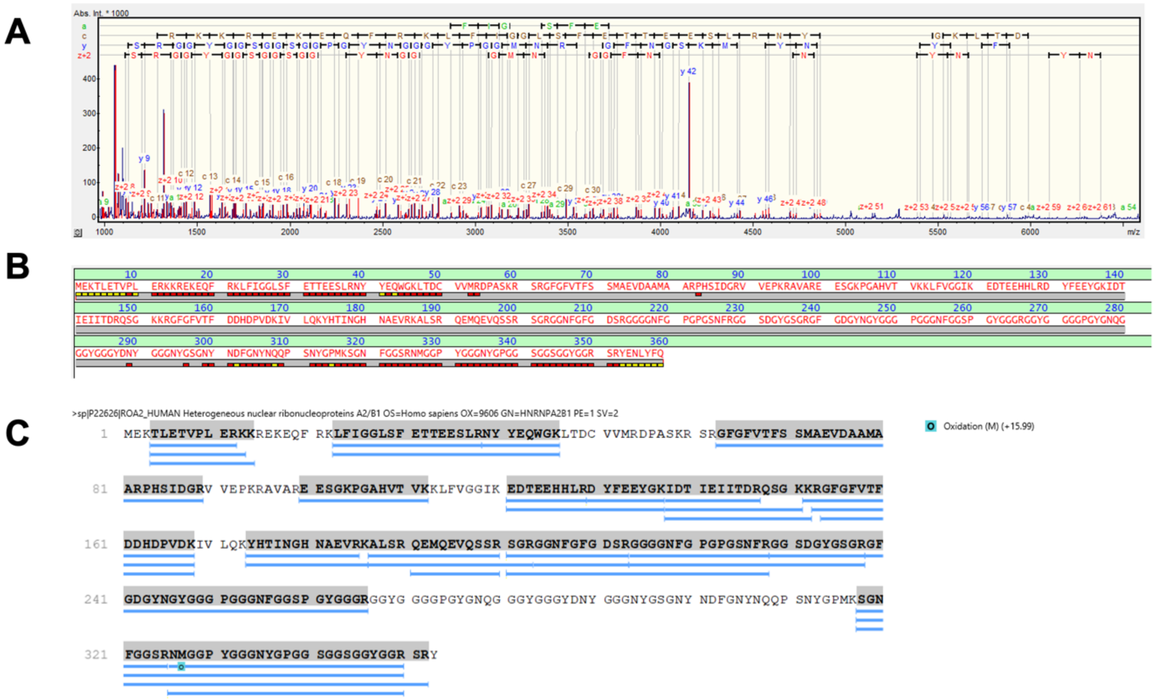


**Figure S.3 Mass Spectrometry analysis of hnRNPA2B1**

*(A) MALDI-ISD-TOF spectrum with all the annotated theoretical fragments of hnRNPA2B1. (B) Top-down sequencing of the N-terminal and C-terminal of hnRNPA2B1. The red blocks indicate matches with the sequence, confirming the expected N- and C-terminal ends. (C) Alignment of the sequence of hnRNPA2B1 with all the identified peptides of a trypsin digest after LC-MS/MS analysis, confirming the identity of hnRNPA2B1. Blue bars represent peptides that were identified based on their molecular weight and sequence.*

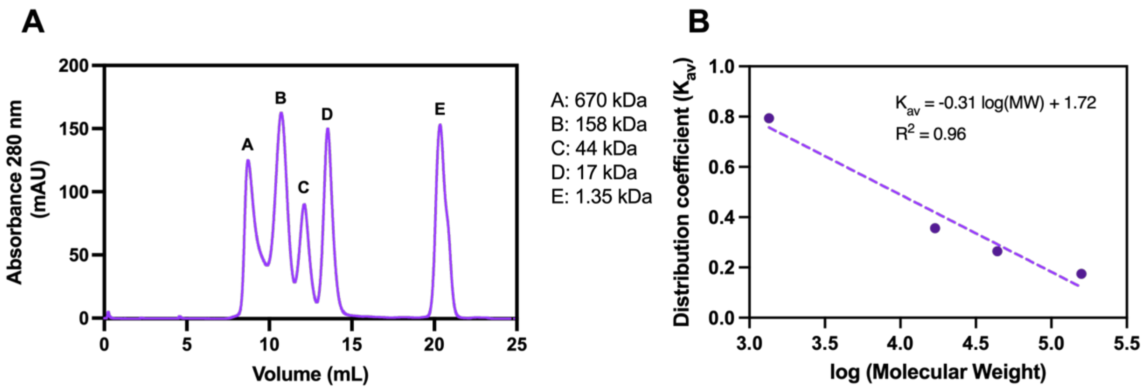


**Figure S.4 Analytical Size Exclusion Chromatography of hnRNPA2B1**

*(A) SEC profile of the Gel Filtration Standards. A: thyroglobulin (670 kDa); B: bovine γ-globulin (158 kDa), C: chicken ovalbumin (44 kDa), D: equine myoglobin (17 kDa), and E: vitamin B12 (1.35 kDa). (B) Calibration curve of the distribution coefficient each standard (expect the thyroglobulin) against the logarithm of the molecular weight.*

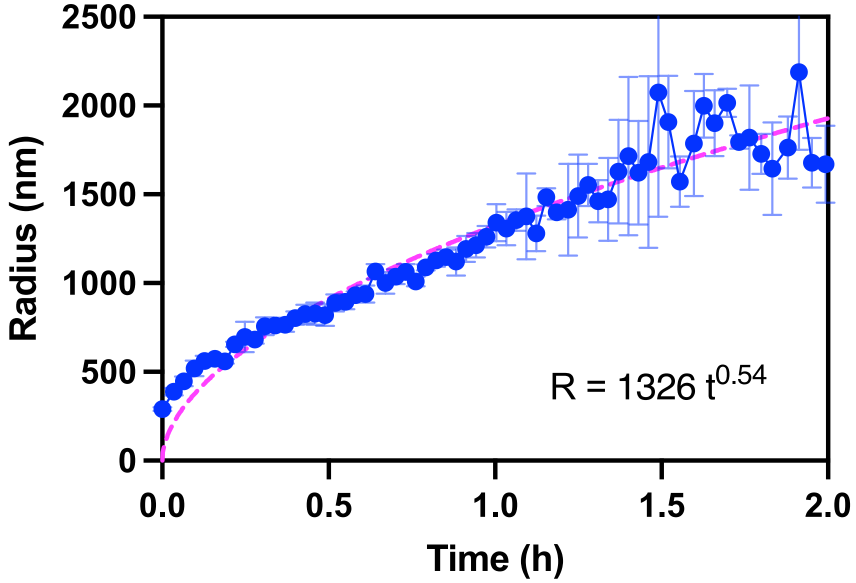


**Figure S.5 Growth kinetics of hnRNPA2B1 droplets measured by DLS.**
*After one hour, the sample became more polydisperse due to the formation of larger droplets and protein aggregation and the data could not be interpreted anymore.*

**Table S.1 NanoDrop Readings.**

| Sample | Protein concentration (mg/mL) | A280 | A260/A280 |
|---|---|---|---|
| HisTrap | 2.78 | 2.20 | 0.59 |
| Desalting | 1.44 | 1.17 | 0.59 |
| Reverse HisTrap | 0.87 | 0.86 | 0.58 |
| Final Dialysis | 0.64 | 0.66 | 1.52 |
